# Large-scale analysis of redox-sensitive conditionally disordered protein regions reveal their widespread nature and key roles in high-level eukaryotic processes

**DOI:** 10.1101/412692

**Authors:** Gábor Erdős, Bálint Mészáros, Dana Reichmann, Zsuzsanna Dosztányi

## Abstract

Recently developed quantitative redox proteomic studies enable the direct identification of redox-sensing cysteine residues that regulate the functional behavior of target proteins in response to changing levels of reactive oxygen species (ROS). At the molecular level, redox regulation can directly modify the active sites of enzymes, although a growing number of examples indicate the importance of an additional underlying mechanism that involves conditionally disordered proteins. These proteins alter their functional behavior by undergoing a disorder-to-order transition in response to changing redox conditions. However, the extent to which this mechanism is used in various proteomes is currently unknown. Here, we use a recently developed sequence-based prediction tool incorporated into the IUPred2A web server to estimate redox-sensitive conditionally disordered regions on a large scale. We show that redox-sensitive conditional disorder is fairly widespread in various proteomes and that its presence strongly correlates with the expansion of specific domains in multicellular organisms that largely rely on extra stability provided by disulfide bonds or zinc ion binding. The analyses of yeast redox proteomes and human disease data further underlie the significance of this phenomenon in the regulation of a wide range of biological processes, as well as its biomedical importance.

## Introduction

The key to the extraordinary versatility of proteins lies in their specific structural properties that precisely suit their individual functions. For example, enzymes typically need to adopt a well-defined conformation that ensures the correct orientation of their active site residues, as required for optimal catalytic activity. In contrast, intrinsically disordered proteins and protein regions (IDPs/IDRs) lack a well-defined structure in isolation, and are best characterized by an ensemble of rapidly interconverting conformations [1]. The specific structural properties of IDPs enable them to carry out a different set of functions, which could not be fulfilled by relatively rigid globular domains [2,3]. As a result, the function of many IDPs directly originates from their high intrinsic flexibility, enabling them to act as linkers or spacers [3,4]. However, there is an emerging understanding that the structural behaviour of IDPs is often context-dependent: a disorder-to-order or order-to-disorder transition can be induced as a result of binding to specific macromolecular partners, undergoing post-translational modifications, or changes in environmental factors such as pH and temperature [5]. Recently, redox conditions have emerged as another important factor that can regulate conditional disorder [6]. The growing number of examples of redox-regulated conditionally disordered proteins indicates that this is an important mechanism triggered in response to various forms of oxidative stress or to naturally occurring changes in redox potential [6,7].

Redox regulation is essential for the survival of all organisms, as cells constantly encounter the transient accumulation of reactive oxygen species (ROS). ROS may be generated exogenously or endogenously due to metabolic activity or inflammation, while subsequent oxidative damage can cause widespread protein unfolding and aggregation. It may also contribute to the pathogenesis of many different diseases, as well as to degenerative processes associated with aging. However, ROS can also serve as cellular signaling molecules and regulate various biological processes such as the immune response, cell proliferation, development, and more [8,9]. The specific effects of ROS are captured in large part through the covalent and reversible modifications of specific cysteine residues, which can in turn modify the structural or functional properties of the redox-sensitive target proteins. Such post-translational modifications are tightly regulated by changes in levels of diverse physiological oxidants and is generally faster than expression of specific regulatory proteins. Over the last decade, numerous examples of such thiol-switch proteins have been uncovered, spanning different functions and regulation modes [6,7,10]. The functions of select proteins were found to be regulated by oxidative modifications of a specific cysteine thiol forming either sulfenic (e.g., in 1-Cys peroxiredoxin) or sulfenamide (e.g., in tyrosine phosphatase 1B, leading to regulation of the Ras signaling pathway). Another more studied mode of activation is through disulfide bond formation by redox-sensitive cysteine residues, typically located in close proximity of each other. These disulfide bridges can either promote disorder-to-order transitions (e.g., chloroplast regulator of the calvin cycle, CP12 [11] and cell growth modulator, granulin Grn-3 [12]) or vice versa (e.g., in the bacterial protein chaperone Hsp33 [13] and mammalian copper chaperone Cox17 [14]) in response to shifts in the cellular redox status [6].

The growing interest in redox regulation has motivated the development of various proteomics approaches that can be used to explore the landscape of thiol redox modifications. These methods usually involve a differential modification of the thiol groups by specific probes which labell reduced thiolate groups. Since the oxidized cysteines do not interact with these probes, the typical workflow involves an initial modification of all reduced thiols in the cell proteome, followed by reduction and then labeling with the second probe which differs in mass from the first. Thus, reduced and oxidized cysteines can be differentially labeled in a ratiometric manner, allowing for quantitative analysis of reversible oxidation modification of thiol groups. One of these approaches is based on the differential labeling by N-Ethylmaleimide (NEM) and *N*-(6-(biotinamido)hexyl)-3′(2′-pyridyldithio)propionamide (biotin-HPDP), which enables the analysis mainly of the reversibly oxidized thiols [15]. In contrast, the OxICAT methodology is based on differential labeling of reduced and oxidized cysteines with isotopically light and heavy forms of the isotope coded affinity tag (ICAT) reagent, derivatives of the iodoacetamide alkylation reagent [16]. The usage of two differential isotopes provides quantification of the degree of oxidation among thousands of specific cysteine residues within the analyzed proteome in a highly precise manner. OxICAT has thus far been applied to proteomes (redoxomes) of diverse organisms under different physiological conditions [17–20].

Using redox proteomics studies, the cellular redox state has thus far been revealed to either directly or indirectly affect a wide range of physiological processes in the cell. However, proteomics studies provide no information about the underlying mechanisms of thiol modifications, nor to what degree the detected thiol modifications correspond to redox-regulated conditional disorder. In order to explore this aspect of redox regulation, we took advantage of a recently introduced new feature of the IUPred method (IUPred2A) that enables the sequence based prediction of regions that are likely to undergo disorder-to-order transitions upon redox changes [21]. IUPred is a robust sequence-based prediction method for protein disorder that uses an energy estimation method [22]. The strength of the method originates from its ability to capture the basic biophysical properties of disordered segments: their inability to form enough stabilizing interactions to adopt a well-defined globular structure. The core of the method is a statistical potential that assigns more favorable scores to amino acid pairs that are observed more frequently in the proximity of each other within the structure of globular proteins, as compared to a background model. In this regard, cysteine residues clearly stand out, as they often form covalent contacts with other cysteines, thus providing large stabilizing contributions either by forming disulfide bridges or coordinating Zn^2+^ and other metal ions. Consequently, the cysteine residues have a strong order promoting character in IUPred. However, thiol modifications of either paired or single cysteine residues may lead to a reversible order-to-disorder transformation, exposing unfolded regions crucial for protein function [6,7]. This regulatory effect of redox conditions on protein structure not only follows the presence of oxidation sensitive cysteine residues, but is also influenced by the presence of disorder-promoting residues in the surrounding sequence. In such cases, cysteine residues may simply be considered small, polar residues akin to serine, located within a sequence region exhibiting distinct features. The prediction of redox-sensitive disordered regions is based on capturing this dual characteristics of cysteines. This novel method opens new doors to explore redox-sensitive conditionally disordered segments in various proteomes.

Here, we apply our method to predict redox-sensitive disordered regions on a large scale. We analyze the occurrence of these segments in known structures, and connect them to structural and functional features. In order to explore evolutionary trends of redox-sensitive conditionally disordered regions, we estimate their abundance at the proteome level and analyze their enrichment in terms of domains and biological processes. By taking advantage of recent mass spectrometry-based techniques applied to the exploration of redox-sensitive thiols, we can highlight proteins that are likely to use disorder-to-order transition to respond to redox changes in biological settings. We have used experimentally derived datasets generated by two different redox probes: either ICAT (used in the OxICAT redox proteomic method) [16] or biotin-HPDP [15]. Finally, by analyzing disease data, we collected examples of genetic mutations that alter the behaviour of putative redox-sensitive regions.

## Results

### I. Prediction of redox-sensitive structural switches in proteins for known examples

Redox-dependent disorder-to-order transitions have been described in detail in only a few cases [6,7]. For this reason, rigorous benchmarking on a large number of positive and negative data is not possible for the prediction of redox-sensitive structural transitions. Instead, IUPred2A was established and is further customized in this study, to best describe the available experimentally verified examples. The prediction is based on the generation of two prediction profiles, one using the original amino acid sequence (redox_plus, assuming cysteine stabilization) and one with mutating each cysteine to serine (redox_minus, corresponding to the lack of cysteine stabilization). While the prediction scores would differ for any sequences containing cysteines, in most cases, no changes in the overall order-disorder tendency would be observed. Our hypothesis was that in the cases of sequences that undergo disorder-to-order transition upon changes in redox conditions, a pronounced shift could be observed with the redox_plus profile predicted as ordered, and the redox_minus profile predicted as disordered. Redox-sensitive regions involved in this type of transition are predicted based on the observed divergence of the two profiles (See Data and Methods).

The performance of our prediction method is illustrated through three interesting cases. One of the best characterized examples include the Hsp33 heat shock protein from *E. coli*. This protein chaperone forms a well-defined structure under reducing conditions with the N-terminal domain in close contact with the adjacent linker region and Zn^2+^ binding C-terminal domain [27]. However, substrate binding is inaccessible in this conformation and consequently, the protein lacks chaperone activity[13]. Oxidative unfolding (oxidation coupled with mild protein destabilization conditions) causes the C-terminal region to release the Zn^2+^ ions, inducing the unfolding of the C-terminal domain together with the adjacent linker region. This transition exposes the substrate binding sites that makes the protein active [28,29]. The critical region for redox-sensing is located between resides 230 and 266, which contains the four highly conserved cysteine residues that coordinate the Zn^2+^ ion under reducing conditions, but form short-range disulfide bonds under oxidative conditions. The large part of the experimentally determined redox-sensitive region is correctly identified by IUPred2A (Figure 1) [30,31].

**Figure 1:**
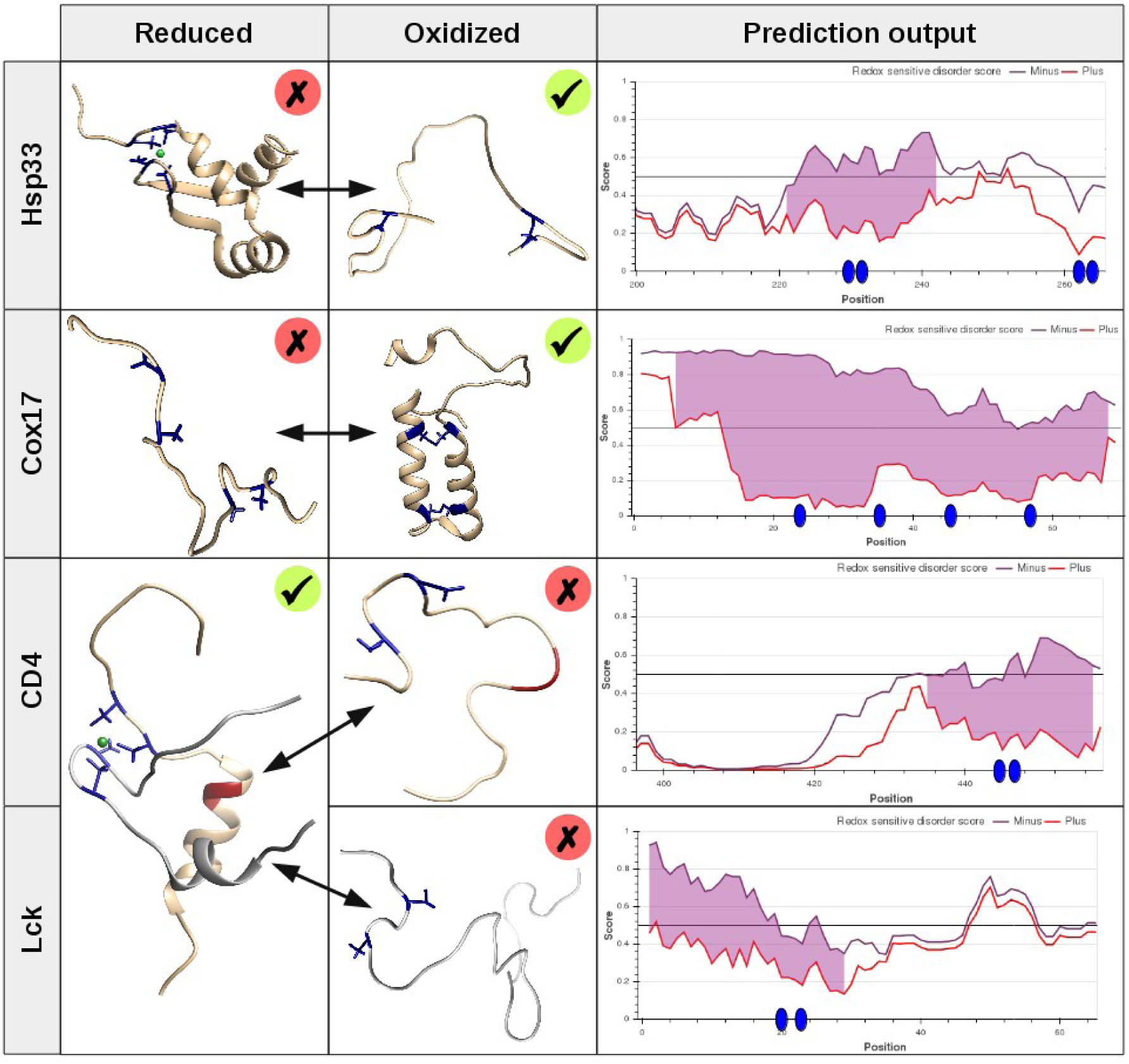
Known examples of redox-change induced structural transitions. Structural figures: cysteines are shown in blue, metal ions are marked as green spheres. The di-leucine degradation motif of CD4 is shown in red. Green ticks and red crosses mark the active and inactive states of the proteins/complexes. The prediction output is taken from the IUPred2A web server, with purple shadings marking the predicted redox-sensitive regions, and blue ovals showing the positions of the redox-sensitive Cys residues. UniProt accessions used: P0A6Y5 (Hsp33), Q12287 (COX17), P01730 (CD4), P06239 (Lck). PDB IDs used: Hsp33|reduced (1xjh), COX17|oxidized (1z2g), CD4:Lck|reduced (1q68). Structures of Hsp33, CD4 and Lck feature only the redox-sensitive regions.

Another prototypical example of redox-regulated disorder-to-order transitions is COX17, a ~60 residue copper chaperone which is one of the 13 subunits of the mammalian cytochrome c oxidase complex. This protein is disordered under the reducing conditions of the cytosol, in which it is synthesized, and its disordered nature is essential for its diffusion across the mitochondrial outer membrane [14,32,33]. However, upon its entry into the oxidizing environment of intermembrane space of the mitochondria, COX17 undergoes a disorder-to-order transition, and its folding proceeds by sequential disulfide formation and association with the Mia40 protein. In agreement with experimental data, the whole protein - apart from its signal sequence - is predicted to undergo a disorder-to-order transition upon changes of redox potential.

The redox-regulated order-to-disorder transition in the case of Hsp33 and the disorder-to-order transition of COX17 produce the active form of the proteins, however, such transitions can also be part of more complex regulatory mechanisms. For example, the cytoplasmic tail of T cell coreceptors CD4 and CD8 associate with the N-terminus of the Src-family tyrosine kinase Lck. These interactions are critical for T cell development and activation. The interacting tail regions of both CD4 and Lck were shown to be intrinsically disordered in isolation, yet assemble to a form a zinc clasp structure [34]. Interestingly, the interaction is also regulated at the level of alternative splicing [35]. The interaction prevents the internalization and degradation of CD4 by masking the dileucine motif required for the clathrin-mediated endocytosis [34]. While there is no direct evidence for redox regulation of this complex, activation and proliferation of CD4+ T cells is associated with the intracellular increase of reactive oxygen species and release of zinc [36]. This suggests that the Lck-CD4 complex could be considered as an additional example of the redox-regulated conditional disorder.

### II. Redox-sensitive conditionally disordered regions in the PDB

The number of fully characterized redox-induced folding/unfolding examples are rather limited. However, the developed method can be further characterized and refined using the vast amount of information encoded in protein structures containing either oxidized or reduced forms of cysteines in the PDB. To this end, we collected structures from the PDB using a filtering of 40% sequence identity. The predictions were run using the sequence from the corresponding PDB file. The presence of disulfide bonds was calculated from the atomic coordinates. However, cysteine residues that are involved in coordinating metal ions are also often located in close proximity. Such clusters were also identified based on the coordinates (see Data and Methods). The statistics for each PDB chain in the non-redundant dataset is given in the supplementary material. Redox-sensitive regions were predicted for 711 out of 14,145 structures, representing only 5.02% of cases (Figure 2A). Most of the redox-sensitive regions corresponded to structures stabilized by disulfide bridges (417, 2.95 %), or cysteine clusters, coordinating either Zn^2+^, Cu^2+^ or Fe^2+^ clusters (214, 1.51%). However, 1,566 structures with disulfide bonds and 344 structures with cysteine clusters did not contain predicted redox-sensitive structural switches. In general, structures with redox-sensitive regions had a higher number of disulfide bonds or cysteine clusters (Figure 2B and 2C).

**Figure 2:**
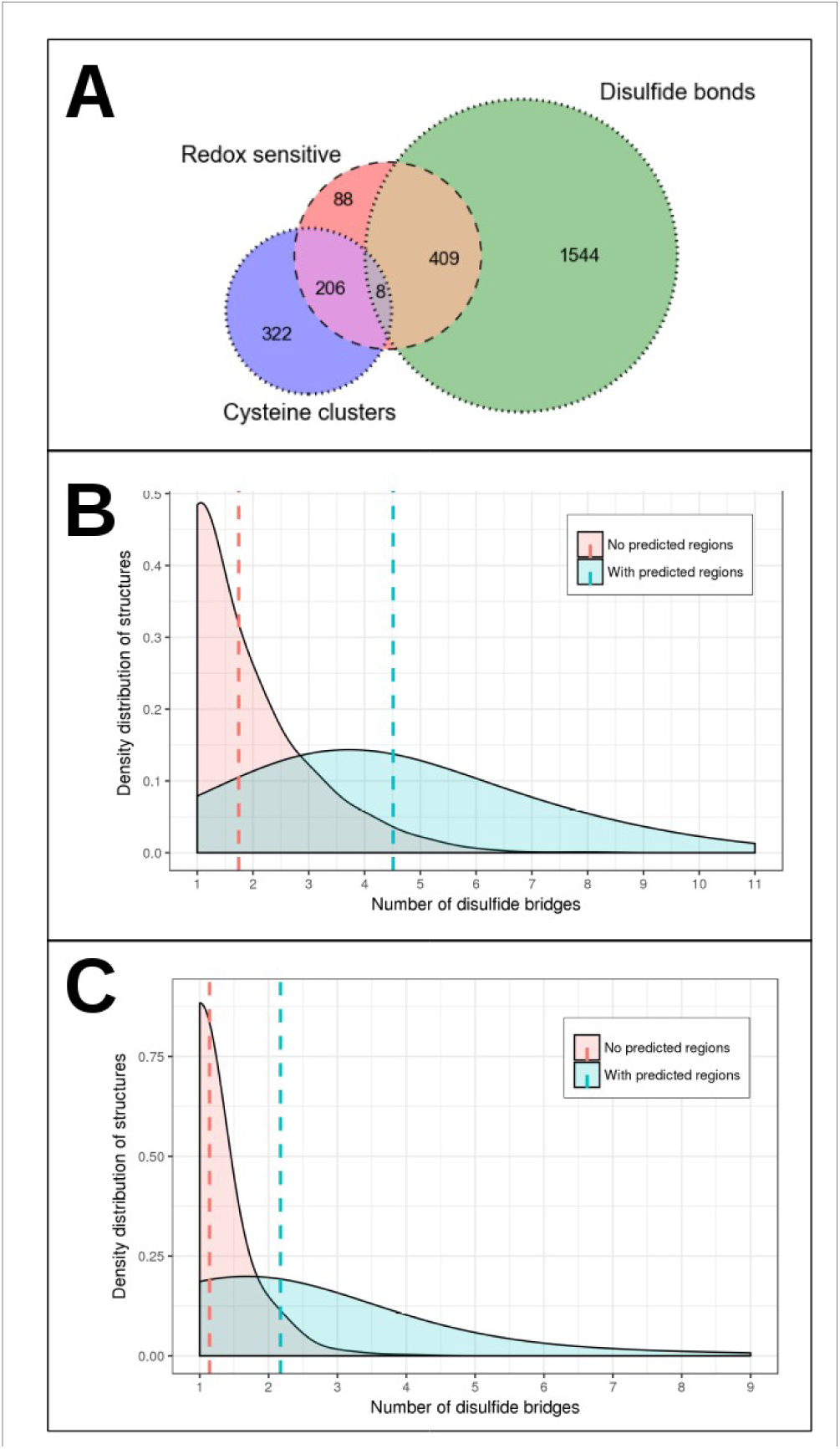
Predicted redox-sensitive regions in the PDB. The overlap between proteins with predicted redox-sensitive regions, and proteins with disulfide bonds/metal ion Cys clusters in the PDB (A). The distribution of structures with respect to disulfide bridge (B) and metal ion coordinating Cys clusters (C) for structures with and without predicted redox-sensitive regions. Dashed lines mark the average values for all distributions.

While disulfide bridges usually from under oxidative conditions, metal binding is more common in reducing environment. Consequently, these structural features are usually complementary and there are only, a few structures contained both cysteine clusters and disulfide bonds. While in certain cases the disulfide bond formation could be a crystallization artefact (e.g.2g45), in many cases, the complementary nature of metal binding and disulfide bond formation can ensure a stable structure under a variety of conditions. As a counterpoint to structures with both of the stabilizing features, a relatively small number of predicted proteins (88, 0.62%) did not have either. The majority of these structures describe heme binding proteins, which rely on heme groups coordinated partially by cysteines for their stability, and might in fact be involved in redox regulation [37]. However, disorder-to-order transitions are not necessarily coupled to redox regulation. In general, examples without the capacity to form either disulfide bonds or cysteine clusters are treated as likely false positives. However, the number of such cases in PDB structures represent less than 1% of all structures in our dataset (Figure 2A). The overall relatively low rate of predicted redox-sensitive conditionally disordered regions in known structures and their strong correlation with strongly stabilizing structural features add strong credibility to IUPred2A redox-sensitive predictions.

### III. Redox-sensitive structural switching correlates with evolutionary complexity

IUPred2A is able to predict redox-sensitive protein regions based on sequences alone. This opens up the means of studying the large-scale distribution of such regions across several proteomes (see Data and Methods), to gain evolutionary insights into the emergence of redox-sensing structural switches. We assessed the fraction of proteins with redox-sensitive regions with a potential structural plasticity in reference proteomes from all three domains of life, as well as viruses (Figure 3A).

**Figure 3:**
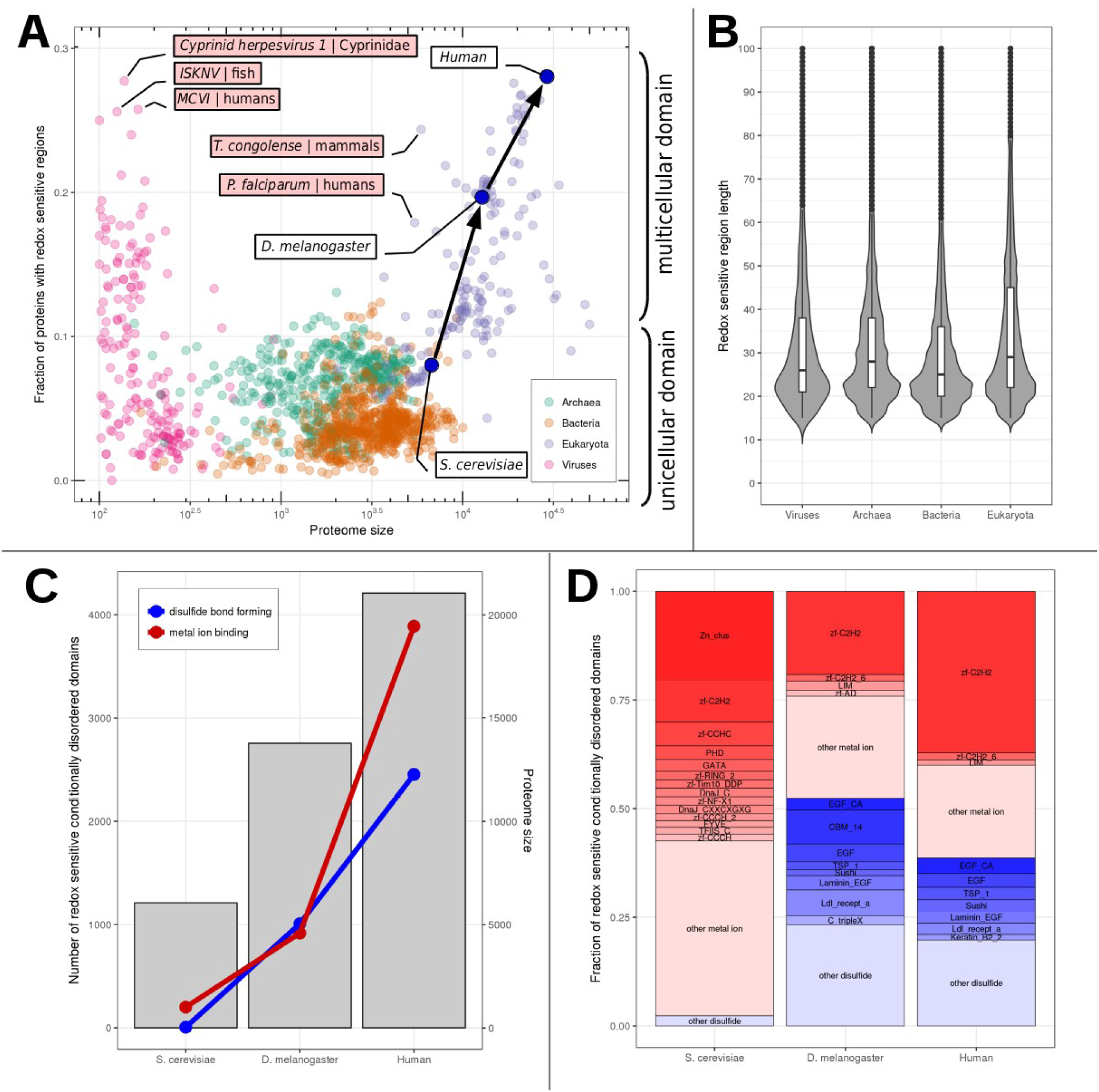
Proteome-wide distribution of redox-sensitive protein regions. A: the fraction of proteins containing redox-sensitive regions as a function of proteome size (on a logarithmic scale). Red boxes mark pathogens, with the host organisms following the name. B: Length distribution of redox-sensitive regions across various domains of life and viruses. C: The number of predicted redox-sensitive metal ion coordinating (red) and disulfide bonded (blue) conditionally disordered structural units (left vertical axis). Proteome sizes are shown in grey bars for reference (right vertical axis). D: The frequency of various Pfam domains among the predicted redox-sensitive structural switches (see supplementary material for full lists). Red boxes mark metal ion coordinating domains, and blue boxes mark disulfide bonded domains. Only domains with a relative frequency of 1.25% or higher are shown in separate boxes, other domain types are merged and shown as “other”.

Genome size generally reflects the average complexity of the domains of life. While the differences between specific organisms are obscured by the C-value paradox, generally viruses have the smallest genomes, followed by archaea, bacteria, and eukaryotes. This hierarchy is also reflected in the proteome sizes, albeit with large overlaps between various domains of life. However, the fraction of predicted redox-sensitive intrinsically disordered proteins (termed f_redox_, see Data and Methods) shows a distinctively different distribution. Bacteria feature the lowest f_redox_ values, with over 90% of species having f_redox_<7%. In general, for unicellular organisms, f_redox_ is below 10%, including almost all archaea and bacteria, together with some eukaryotes. In contrast, multicellular organisms are characterized by markedly higher f_redox_ values, with some highly complex organisms reaching close to 30%. The steady increase of f_redox_ with the increase of complexity is demonstrated by the comparison of the *S. cerevisiae* (f_redox_=7.2%), *D. melanogaster* (f_redox_=19.6%) and human proteomes (f_redox_=27.2%). This highlights that f_redox_ values (as opposed to genome or proteome sizes) better reflect the apparent complexity of organisms.

For complex organisms, the derivation of even the proteome size from the genome sequence is non-trivial[38]. This is further modulated by the presence of various isoforms; therefore in theory, f_redox_ values could be highly dependent on the exact definition of the proteome used for calculations. However, testing for this effect showed that in reality the calculated f_redox_ values are strikingly stable with respect to such variations in proteome definitions. The minimal proteome of *D. melanogaster* from UniProt contains 3,496 proteins, while the proteome featuring all isoforms contains 13,775 proteins. Despite this nearly 4-fold difference, both proteomes yield almost identical f_redox_ values of 19.6% and 20.8%, respectively. This supports the robustness of f_redox_ as a marker for organism complexity, with the largest breakpoint corresponding to the advent of multicellularity.

Apart from multicellular eukaryotes, several viruses also exhibit high f_redox_ values. In these cases, f_redox_ values correlate with the complexity of the host, with most high f_redox_ value viruses targeting complex eukaryotes, typically vertebrates. This correlation between host complexity and f_redox_ value is also apparent for several unicellular eukaryotic pathogens. Both *T. congolense* and *P. falciparum* (the major causes of the sleeping sickness and malaria, respectively) are unicellular eukaryotes, and such their f_redox_ values are expected to be below 10%, similarly to other organisms with comparable complexity and small proteome size. However, their f_redox_ values rather resemble those of their hosts, with f_redox_=24.4% and 17.9%, respectively (Figure 3A).

While the fraction of proteins incorporating redox-sensitive intrinsically disordered regions increases with organism complexity, the typical length of such regions is strikingly constant across various domains of life and viruses (Figure 3B). The typical region length is approximately 22 residues, and while eukaryotes have a slightly higher average region length, this is primarily a result of the high apparent length of tandem arrays of redox-sensing regions. This indicates that the basic molecular mechanisms behind redox-sensing structural switches are probably uniform across all life, and the emergence of complexity requires a larger number of proteins utilizing such mechanisms.

In order to directly assess the structural units involved in redox-sensitive order-disorder transitions, we also analyzed the domains that overlap with the predicted regions in the three model organisms highlighted in Figure 3A (see Figure 3C). In the yeast proteome, nearly all redox-sensitive structural switches are metal ion coordinating domains. The majority of these domains coordinate zinc ions, belonging to a handful of zinc finger domain types (Figure 3D). The transition into the realm of multicellular organisms brings about a drastic increase in the relative number of structural switches harboring zinc ion coordinating structural switches. While the ratio of proteome sizes for yeast vs. drosophila is only 2.3 (6,049 vs. 13,775), the ratio of the number of identified metal ion coordinating domains is about 4.5. Analysis of the drosophila proteome also demonstrates the emerging dominance of C2H2-type zinc fingers. In addition, the drosophila proteome also shows a striking increase in the number of disulfide-bound structural switches. In contrast to metal ion binding domains, however, the identified disulfide bonded domains are more evenly distributed across several types of domains. Considering the human proteome, the abundance of both types of structural switches are even higher. However, this increase is more pronounced for the metal ion binding domains, and is largely attributable to the drastic increase in the number of C2H2 type zinc finger domains. In comparison, extracellular, disulfide bonded domains remain fairly diversified across several domain types.

Judging by sheer numbers of occurrence, the most abundant domain of interest in humans is the C2H2 zinc finger. This structural unit is present in all three studied proteomes in high numbers. Furthermore, the ratio of conditionally disordered C2H2 domains compared to the total number of such domains is surprisingly stable across the three organisms, with values of 59.5%, 66.6%, and 53.9%, for yeast, drosophila and human. This indicates that the study of the redox-sensitivity of simple model organisms, such as yeast can have implications in human physiology as well. The identification and functional characterization of redox-sensing structural switches can therefore serve as a guide to the understanding of the more complex roles these protein regions play in human regulation.

### IV. Comparison with redox proteomics datasets in yeast

We analyzed three datasets of *S. cerevisiae* proteins containing modified cysteines identified using redox proteomics. In the first case, 27 proteins were identified harboring reversibly-oxidized cysteines using biotin-HPDP-based redox proteomics[15]. The biotin-HPDP-based method identifies only cysteines that were oxidized in the protein lysate. On the other hand, the other two datasets were obtained using the OxICAT method, which analyses all cysteines, and provides degree of oxidation of the identified cysteines (reduced and oxidized in the protein lysate), focusing on a wider range of cysteine containing peptides. The second study applied quantitative redox proteomics based on the OxICAT approach[39] and identified 41 proteins that underwent substantial thiol modifications as a result of H_2_O_2_ treatment. Using similar thiol labeling but different mass spectrometry workflow, the third study identified 47 unique proteins with redox-active thiols[40]. Strikingly, there was very limited overlap between these datasets. Only five proteins were common between the first two studies (TRP5, TEF1, FBA1, ARO4 and AHP1), and only a single protein (RPL37B) was shared between the second and third studies, with no overlap between the first and third studies. This small overlap among the three studies seems to indicate limitations in terms of the currently applied techniques’ sensitivity, as well as technical and biological variabilities.

The structural/functional consequences of cysteine modifications identified in these experiments can be heterogeneous, and not all of them are expected to induce a transition in the structural state of the affected region. Nevertheless, in several cases, the identified proteins with modified cysteines overlapped with predicted conditionally disordered regions. As these predictions suggest a specific mechanism underlying the redox-sensitivity, we took a closer look at these examples (Table 1). In one example, the GAS1 protein is anchored to the plasma membrane and to the cell wall. In addition to the catalytic glucanosyltransferase domain, it also contains a cysteine rich domain. The redox-sensitivity of these cysteines in not easy to explain, as they are required for the normal folding and stability of the protein[41]. An interesting case corresponds to YDJ1, a yeast homolog of the chaperone DnaJ, a zinc containing co-chaperone, involved in mitochondrial protein import. The central cysteine-rich domain, which contains four repeats of the motif CXXCXGXG, is predicted as conditionally disordered. Its redox-sensitivity suggests that it could be involved in oxidative-stress response, similarly to Hsp33. The validity of the prediction is supported by the notion that redox regulation was established for the human homologue of DnaJ [42].

**Table 1.** Experimentally characterized redox-sensitive yeast proteins containing predicted conditionally disordered regions.

| Uniprot accession | Gene name | Protein name | Redox cysteines | Predicted region | Affected Pfam domain | Reference |
| --- | --- | --- | --- | --- | --- | --- |
| P22146 | GAS1 | 1,3-beta-glucanosyltransferase GAS1 | 421 | 360-385 | X8 (disulfide bond) | [15] |
| P25491 | YDJ1 | Mitochondrial protein import protein MAS5 | 185, 188 | 135-213 | DnaJ_C (Zn<sup>2+</sup> coordinating) | [39] |
| P41058 | RPS29A, RPS29B | 40S ribosomal protein S29-B | 24, 39 | 11-31 | Ribosomal_S14 (Zn<sup>2+</sup> coordinating) | [40] |
| P0CH09 | RPL40A, RPL40B | Ubiquitin-60S ribosomal protein L40 | 115 | 82-124 | Ribosomal_L40e (Zn<sup>2+</sup> coordinating) | [40] |
| P51402 | RPL37A, RPL37B | 60S ribosomal protein L37-B | 19, 34, 37 | 9-46 | Ribosomal_L37e (Zn<sup>2+</sup> coordinating) | [39,40] |
| P25348 | MRPL32 | 54S ribosomal protein L32, mitochondrial | 104, 107 | 97-116 | Ribosomal_L32p (Zn<sup>2+</sup> coordinating) | [40] |
| P10962 | MAK16 | Protein MAK16 | 38 | 18-51 | Ribosomal_L28e (Zn<sup>2+</sup> coordinating) | [40] |
| P36053 | ELF1 | Transcription elongation factor 1 | 25, 28, 52 | 15-61 | Elf1 (Zn<sup>2+</sup> coordinating) | [40] |
| P46677 | TAF1 | Transcription initiation factor TFIID subunit 1 | 1038,1041 | 1031-1053 | zf-CCHC_6 (Zn<sup>2+</sup> coordinating) | [40] |
| P53849 | GIS2 | Zinc finger protein GIS2 | 107 | 1-152 | zf-CCHC (Zn<sup>2+</sup> coordinating) | [40] |
| P32770 | NRP1 | Asparagine-rich protein | 378 | 351-387 | zf-RanBP (Zn<sup>2+</sup> coordinating) | [40] |
| P39956 | RPH1 | DNA damage-responsive transcriptional repressor RPH1 | 740 | 733-772 | zf-C2H2 (Zn<sup>2+</sup> coordinating) | [40] |
| P40422 | RPC10 | DNA-directed RNA polymerases I, II, and III subunit RPABC4 | 31, 34 | 38-59 | DNA_RNApol_7k D (Zn<sup>2+</sup> coordinating) | [40] |
| Q06835 | RDS3 | Pre-mRNA-splicing factor RDS3 | 23 | 17-59 | PHF5 (Zn<sup>2+</sup> coordinating) | [40] |
| Q02772 | PET191 | Mitochondrial protein PET191 | 32 | 1-55 | Pet191_N (disulfide bond) | [40] |

Several members of the ribosome were also shown to be redox-sensitive, further containing predicted conditionally disordered regions. This group includes the mitochondrial Mrpl32, which was shown to undergo redox-dependent regulation[43]. Members of the cytosolic ribonucleoprotein complex were also included in this group (RPL37A, RPL37B, RPS29A, RPS29B, RPL40A, RPL40B). In addition, MAK16 is involved in the biogenesis of the ribosomal large subunit in the nucleolus. The common structural theme among these proteins is Zn^2+^ binding, typically involving a specific zinc finger class, called treble clefs. Additional members of the translation machinery are also candidates for redox-regulated conditional disorder mediated by their Zn^2+^ binding domains, such as Taf1 and Elf1. Taf1 is a zinc knuckle that plays a central role in TFIID promoter binding and regulation of transcription initiation[44]. Elf1 is a conserved transcription elongation factor that contains a putative zinc binding domain with four conserved cysteines[45]. Another Zn^2+^ binding protein, Gis2 interacts with the mRNA translation machinery and is a component of the stress induced RNP granules [46]. NRP1 also belongs to the group of Zn^2+^ finger proteins, with a currently uncharacterized RNA binding function, also localizing to stress granules. From a functional point of view, the redox-sensitivity of these proteins can play a critical role in the global attenuation of translation in response to oxidative stress [40]. Further candidates for redox-regulated conditional disorder include other Zn^2+^ binding proteins, such as the DNA damage-responsive transcriptional repressor RPH1, Pre-mRNA-splicing factor RDS3 and DNA-directed RNA polymerases I, II, and III subunit RPABC4 (Table 1).

In contrast to the other examples, redox-sensitive cysteines in Pet191 are likely to form disulfide bonds when it is located in the intermembrane space of the mitochondria. This protein is a cytochrome c oxidase assembly factor with a similar twin-Cx_9_C motif, similar to that found in Cox17. The disulfide bond formation is required for cell viability and assembly of the cytochrome c oxidase complex, however the protein may be imported into the mitochondria, independently of the mitochondrial MIA40-ERV1 machinery [47]. This difference could explain why this protein was found to be redox-sensitive, while other members of the COX family were not detected as such.

### V. Functional repertoire of redox-sensitive structural switches in the human proteome

Yeast proteomics results highlight some key molecular mechanisms in which redox-sensitive conditional disorder plays functional roles, most notably in stress response, biogenesis, and the regulation of transcription and translation. These results indicate that these structural switches preferentially contribute to the actuation of certain biological processes. While yeast proteomics can highlight certain functions also present in higher order organisms, biological processes that are unique to higher order multicellular organisms can only be uncovered via the direct analysis of the functional annotations of their proteomes.

The characteristic functional repertoire of human redox-sensitive conditionally disordered proteins was assessed using Gene Ontology (GO) term enrichment analysis. We identified terms that are significantly enriched in the set of predicted conditionally disordered proteins compared to the full human proteome (see Data and Methods). Relevant GO terms were divided into four groups, termed slims, to separately highlight basic molecular (GO molecular slim), network/pathway level (GO network slim), cellular level (GO cell slim), and organism level biological processes (GO organism slim).

Molecular level functions highlight that the dominant processes uncovered by yeast proteomics are also heavily associated with redox-sensitive conditional disorder in the human proteome. These processes are linked to the regulation of transcription and translation, and are dominated by intracellular proteins. Consequently, the corresponding GO terms are clearly linked to metal ion coordinating regions. In addition, these structural elements are also associated with the biosynthesis of several organic compounds, processes that are largely absent from the results of yeast proteomics. While corresponding processes do exist in yeast, they involve only a very limited number of proteins in contrast to the similar function in humans (e.g. ‘*nucleobase-containing compound metabolic process*’ covering only 7 yeast proteins [48]). A separate class of GO molecular slim terms correspond to the multicellular-specific metabolism and organization of the extracellular matrix in general, and collagen fibrils in particular. These processes are clearly dominated by extracellular proteins, and are thus strongly associated with redox-sensitive disulfide bridge forming protein regions.

Analysis of network level processes shows that redox-sensitive conditionally disordered proteins are heavily involved in the regulation of key processes. Cell growth is governed by several signals, and the processing of various growth factors (such as TGFbeta) and retinoic acid is clearly dependent on conditional disorder, with a slight preference for disulfide bonded proteins. In contrast, steroid hormone signal processing mainly involves intracellular, metal ion coordinating structural switches. Generic cell-cell communication involving various members of the integrin receptor family relies on a balanced mix of disulfide bonded and metal ion coordinating structural elements. A clear exception to this rule is the interaction between nerve cells, as the assembly and organization of synapses very heavily utilizes extracellular conditionally disordered proteins, stabilized by disulfide bonds.

These molecular and network level processes enable characteristic cellular functions that are typically centered around cell movement, growth, differentiation, and proliferation. The majority of these processes require the coordination between various pathways and utilize several molecular processes. As a result, these processes typically show very little preference for either of the two types of redox-sensitive structural switches. The two main exceptions to this rule are cell movement via chemotaxis, which mostly involves mostly disulfide bonded extracellular proteins, and stem cell division and population maintenance, mostly relying on metal ion coordinating cysteine clusters.

At the organism level, the cellular functions involving redox-regulated conditionally disordered protein regions are typically embedded in a wide variety of developmental processes. Several of these processes are focused on either reproduction, such as the formation of a labyrinthine layer, or the very early developmental stages of life, such as somitogenesis, gastrulation, or endoderm formation. As these high level organismal processes typically rely on a large number of proteins, they do not show exclusivity towards either structural element.

### VI. Redox-sensitive structural switches in disease related proteins

As the identified human redox-sensitive proteins fulfill central biological roles, their modulation by mutations is expected to have serious physiological consequences. We used the Humsavar dataset of disease associated germline mutations collected in Uniprot[49] to assess human pathogenic conditions associated with the perturbation of redox-sensitive conditionally disordered regions. We collected disease mutations that either eliminate or introduce cysteine residues in the sequence. In general, cysteine is one of the least common amino acids in proteins. Accordingly, the frequency of cysteine residue in sequences with diseases mutations is only 2.44%. However, its frequency among mutated residues is 6.85%, which places it as the second most mutation-prone residue (after arginine) in the disease dataset. We analyzed these cases to establish a link between the induced structural changes, protein function, and phenotypic alterations.

The structural effect of cysteine modifying mutations shows a clear correlation with the subcellular localization of the affected protein. Extracellular regions are targeted through cysteines involved in the formation of the stabilizing disulfide bond patterns, while intracellular proteins are targeted through modulation of metal ion binding (Table 2). Extracellular disulfide patterns can be disrupted by both the introduction and elimination of a cysteine, as both lead to an uneven number of cysteines, resulting in incorrect intra- or intermolecular bond formation. The native disulfide patterns are typically highly conserved [50] and apart from their direct enthalpic contribution, they also contribute to protein stability and folding through indirect effects[51]. Mutations of such cysteines typically reduce the stability of the implicated domains, resulting in incorrect folding, or promoting aggregation, which affects several human plasma membrane receptors and extracellular proteins.

**Table 2:**
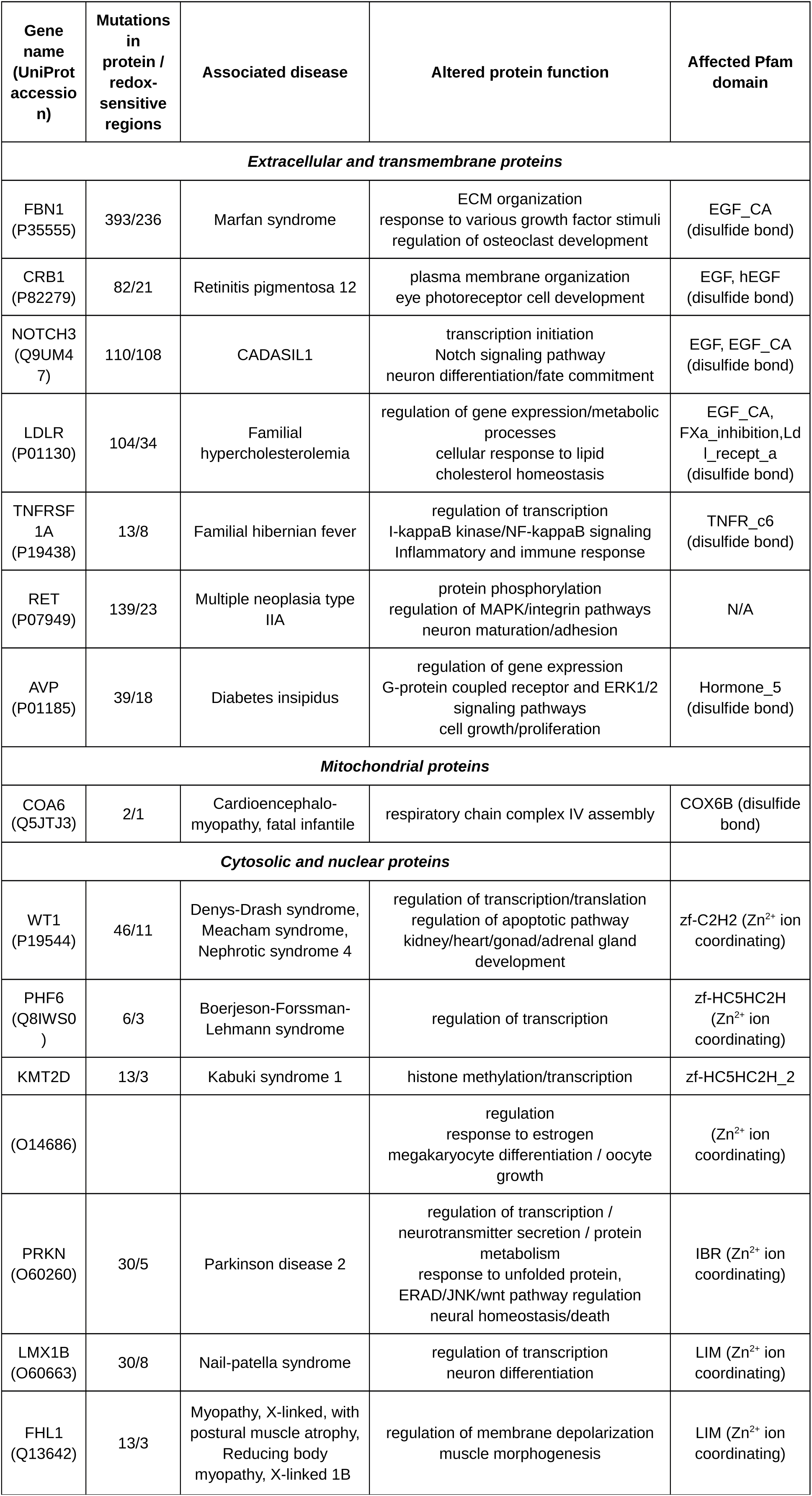
Redox-sensitive regions affected by disease causing germline mutations. For each protein the main disease-affected molecular processes, affected signaling pathways/networks, and cellular/organismal functions are given based on associated GO terms, if available.

One of the most targeted extracellular structural modules in the analyzed dataset are EGF-like domains, which are stabilized via three pairs of cysteines forming three disulfide bonds. EGF-like domains are commonly found in tandem repeats in the ligand binding regions of receptors, and are often targeted by germline mutations. Cys-altering mutations in the EGF domains of **fibrillin**, the major structural constituent of extracellular microfibrils[52,53] results in connective tissue disorders such as the Marfan syndrome. The perturbation results in short or long-range structural rearrangements, and can affect the Ca^2+^ binding ability of the domain, which can lead to increased susceptibility to proteolysis, retention in the endoplasmic reticulum, and delayed secretion[52]. It has been suggested that increased oxidative stress can contribute to disease progression[54]. A radically different phenotype is produced by EGF Cys mutations of **CRB1**, linked to various retinal dystrophies. CRB1 has an important role in maintaining retinal integrity, and its mutations result in ROS accumulation and photoreceptor cell death[55]. Similar mutations in **NOTCH3** cause the vascular dementia disease Cerebral Autosomal Dominant Arteriopathy with Subcortical Infarcts and Leukoencephalopathy (CADASIL). In this case, however, mutations lead to impaired intracellular trafficking and maturation of the mutated Notch 3 receptors[56], and are not directly related to oxidative stress[57]. In NOTCH3, the uneven number of cysteines likely contributes to intermolecular disulfide bridges, as these mutations increase multimerization and aggregate formation [58].

Apart from EGF-like domains, disulfide pattern perturbation by uneven numbers of cysteines was also linked to several other extracellular domains in tandem arrangements. The low-density lipoprotein receptor (**LDLR**) contains a tandem array of LDL-receptor domains and EGF-like domains. LDL-receptor domains are responsible for ligand binding, and EGF-like domains are involved in the degradation and recycling of the receptor. Both types of structural modules are common sites for Cys altering mutations, causing familial hypercholesterolemia[59]. Tumor necrosis factor receptor superfamily member 1A **(TNFRSF1A)** does not contain EGF-like domains, but its extracellular region is composed of four repeats of the TNFR domain, each stabilized by three disulfide bonds. TNRF can harbor Cys-abolishing mutations, disrupting highly conserved intrachain disulfide bonds, leading to the autoimmune disease periodic syndrome phenotypes. This perturbation could contribute to ligand-independent and enhanced ligand-dependent tumor necrosis factor receptor signaling under oxidative stress[60], although impaired cytokine receptor clearance was also proposed as a disease mechanism[61].

Disulfide bond pattern perturbation can also affect individual extracellular domains. The **Ret** receptor tyrosine kinase receptor is mutated in the extracellular juxtamembrane cysteine-rich domain. Similarly to NOTCH3, these mutations induce intermolecular interactions between receptor molecules. The increased dimerization of Ret proteins leads to increased levels of autophosphorylation and tyrosine kinase activity[62,63], underlying a frequent form of medullary thyroid cancer (Ret-MEN2A). Furthermore, cysteine modulating mutations can also severely affect single domain extracellular proteins. A prime example is the arginine vasopressin-neurophysin II (**AVP**-NPII), which is composed of a single domain, stabilized by a network of seven disulfide bonds. Both the creation or the abolishment of a Cys residue are likely to alter the three-dimensional structure of the prohormone, which accumulates in the cell body, ultimately leading to neuronal degeneration and hormonal deficit[64], resulting in autosomal dominant familial neurohypophyseal diabetes insipidus (adFNDI).

The disulfide-favoring oxidizing conditions are not only present in the extracellular space, but can also be found in the intermembrane space of the mitochondria. In the light of our knowledge of redox-sensitive conditionally disordered regions located in this compartment, **COA6** provides an interesting case as regards disease mutations. The protein is involved in the maturation of the mitochondrial respiratory chain complex, and is critical for biogenesis of mtDNA-encoded COX2. A pathogenic mutation in COA6, leading to substitution of a conserved tryptophan for a cysteine residue, results in a loss of complex IV activity and cardiomyopathy. Towards understanding the molecular basis of pathogenesis, it was recently shown that the human COA6 (p.W59C) mutant leads to an increased aggregation state or mislocalization to the mitochondrial matrix[65,66]. In agreement with these observations, the observed mutations introduce a redox-sensitive region, which is not predicted in the native protein sequence.

In contrast to their extracellular counterparts, disease mutations affecting putative redox-sensitive conditionally disordered regions located in proteins inside the cell target metal ion coordinating domains. The majority of such mutations perturb the Zn^2+^ ion binding of various types of zinc finger (ZNF) domains, and in these cases the mutations almost ubiquitously eliminate cysteines. As one of the most common function of ZNF domains is DNA binding, several cysteine-modifying germline mutations affect transcription factors. One of the most well studied such protein is the human zinc-finger transcription factor **WT1**. WT1 ZNF cysteine mutations result in abnormal development of the genitourinary system and are associated with various diseases including Wilms tumors, Denys-Drash syndrome, Nephrotic syndrome 4 and Meacham syndrome. The four C-terminal tandem ZNFs of WT1[67] are mutational hotspots, with the majority of the contained mutations altering the zinc coordinating cysteine residues of ZF2 and ZF3 domains. These mutations can increase the flexibility of the protein or alter its DNA-binding specificity[68]. While there is no direct evidence that WT1 is redox-regulated, in the case of the similar protein EGR1 it was shown that the redox state modules the DNA binding activity of the protein[69,70]. Another transcription factor, **PHF6** is involved in chromatin regulation and neural development[71]. Germline mutations connected to the X-linked mental retardation disorder Börjeson-Forssman-Lehmann syndrome eliminate the conserved C45, C99 and C305 cysteines in either the C2HC or the PHD-type zinc finger. These mutations affect the structure core or zinc coordination of the PHF6-ePHD2 domain and destabilize the correct protein fold and consequently interfere with the normal biological function the protein[71]. Apart from direct DNA binding, ZNF targeting mutations can affect transcription in a more indirect way. C1430R and C1471Y mutations in **KMT2D** likely disrupt the PHD5 finger fold, and reduce histone binding and catalytic activity of the protein[72], leading to Kabuki syndrome.

The functional heterogeneity of ZNF domains results in the germline modulation of redox-sensitive cysteines in proteins with distinctively different biological functions as well. Mutations in Parkin (**PARK2**), a ubiquitin ligase, are one of the predominant hereditary factors underlying some forms of Parkinson disease. Oxidative stress has been recognized as a major contributing factor to the disease[73]. At the protein level, a large fraction of the observed loss-of-function mutations in the hereditary form of the disease destabilize the protein by affecting folding and stability of the complicated Zn^2+^-bound architecture[74,75], leading to impaired degradation of target proteins, such as ZNF746 and BCL2 <u>[76] [76,77]</u>. Other proteins with redox-sensitive cysteine mutations include **FHL1** and **LMX1B**, both containing multiple LIM domains, which are typically involved in protein-protein interactions as opposed to DNA-binding. LIM domains are generally 50–60 amino acids in size and share two characteristic zinc fingers, which are separated by two amino acids[78,79]. LIM domain containing proteins have diverse biological functions, often shuttling between the nucleus and the cytoplasm, potentially even in in a redox-regulated manner[80], acting as adaptors or scaffolds to support the assembly of multimeric protein complexes, and can also contribute to the regulation of the localization[78]. Specifically, LMX1B controls the expression of key genes involved in mitochondrial functions[81]. The FHL1 gene is predominantly expressed in skeletal and cardiac muscle and its mutations are causative for several types of hereditary myopathies. The Cys mutations are likely to significantly affect the structure and the folding of the proteins and have been shown to induce the formation of aggresome-like inclusions[82].

The detailed examples are all modulated through either of the two main redox-sensitive structural elements: disulfide bridges or metal ion clusters. However, the biological functions of these proteins vary, and as a consequence, their mutations lead to very heterogeneous phenotypic alterations. These alterations are enabled by the distinct biological processes and pathways in which these proteins are embedded. Table 2 also shows the relevant molecular, pathway/network, and cellular level processes of the mutated proteins. In some cases the biological processes are not entirely clear, but the majority of the proteins do reflect the general trends of processes utilizing redox-sensitive structural switches (Figure 4).

**Figure 4:**
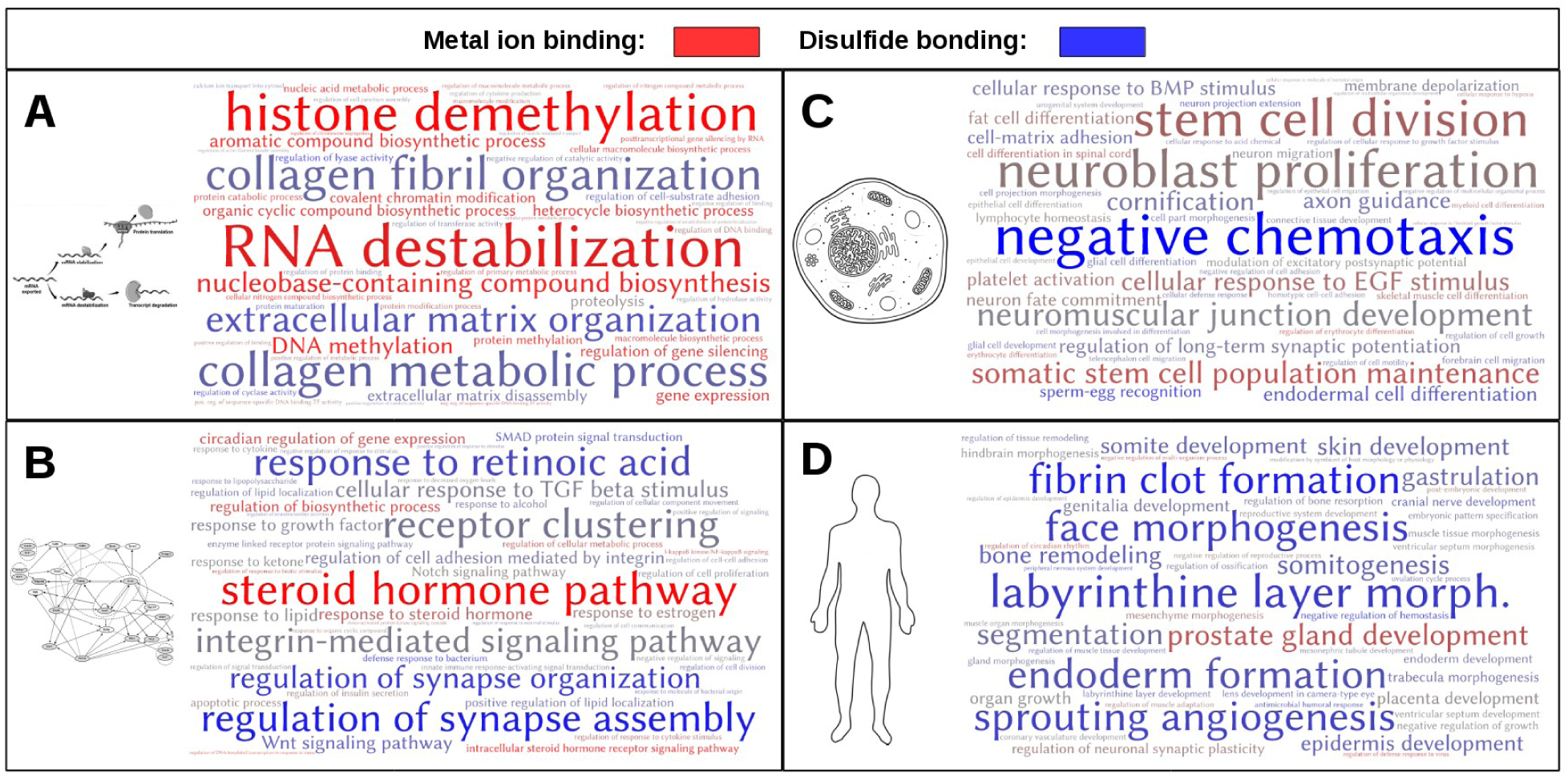
GO term enrichment of human redox-sensitive conditionally disordered proteins. Panels A-D show the 50 GO biological process terms that are most enriched, compared to the full human proteome. Colours denote the dominance of either metal ion coordinating, or disulfide bonded structural elements. Terms that are associated with both structural elements with equal frequency are shown in grey. A: molecular processes, B: pathway/network level processes, C: cellular processes, D: organism level processes. For members of all four slims, together with exact numbers of occurrences and enrichments, see Supplementary material.

## Discussion

Cells constantly encounter changes in oxidative conditions which could be a source of signaling information or can evoke significant stress for the organisms. In order to adequately respond, cells have developed various mechanisms that potentially affect a wide range of physiological processes. Many of these responses are centered around the unique chemical and physical properties of cysteine[83]. Due to its tunable reactivity, cysteine residues are exploited in several different types of proteins including enzymes, signaling proteins, chaperones, and others for regulating their activity. A wide repertoire of potential reversible post-translational modifications of thiol groups enables diverse modes of functional regulation and response to a wide spectrum of physiological oxidants and conditions. Another important property of this residue is its high affinity for metal binding and ability to form disulfide bonds, which provide important additional stabilization for different types of protein structures. As the various functions are often dependent on oxidative conditions, cysteine residues can serve as redox-regulated cellular switches modulating the biological activity of various proteins. It has been demonstrated in the case of a growing number of examples the such thiol switches may trigger conformational changes in proteins, and undergo a disorder-to-order (or order-to-disorder) transition upon changes in cellular redox conditions. Unfortunately, a discovery of such thiol switches encoded in conditionally disordered proteins in the large-scale level is challenging and time-consuming. Therefore, a bioinformatic tool which points to the potential redox-regulated intrinsically disordered region can significantly speed up a discovery of such important regions in a proteome level, pointing on potentially interesting targets for the further biochemical characterization. Here, we have developed and applied such a tool implemented in the IUPred2A web server<u>[21]</u> to uncover potential redox-regulated conditionally disordered protein regions at the proteome level. This opens up a way to assess the functional link between redox-regulated processes and the structural dynamics of the proteins involved.

One of the main outcomes of the presented study is that redox-sensitive conditionally disordered regions (RSCDRs) are widespread in various proteomes, especially in eukaryotic multicellular organisms. According to our predictions, around 5% of the proteome corresponds to proteins with RSCDRs in yeast, however, this proportion increases to nearly 30% in the human proteome. This highly unexpected result gains more credence with the domain-level analysis that established a strong correlation between the increased amount of RSCDRs and the expansion of potentially redox-sensitive domains, such as intracellular zinc fingers or the extracellular EGF domains, at various points of evolution. In yeast, the most common domains predicted to be redox-regulated through conditional disorder are various types of zinc fingers including the DNAJ_C family, or certain subunits of the ribosome. We have used our bioinformatic tool to identify potentially conditionally disordered regions associated with known peroxide-sensitive cysteines in yeast, and identified 15 potentially interesting proteins harboring the RSCDRs (Table 1). This subset of proteins includes ribosomal proteins, a mitochondrial co-chaperone, YDJ1, and transcriptional regulators TAF1, ELF and RDS3. This analysis supports a very-well established interplay between protein and redox homeostasis in cells (29351842), revealing potential members of the protein homeostasis system that might employ redox-regulated conformational changes to regulate their function during oxidative stress. These predictions might direct further experimental analysis of a crosstalk between redox regulation, structural changes, and proteostasis related function. Interestingly, proteins with the predicted redox-regulated disordered regions are predominant in drosophila species. This increase can be partially attributed to the expansion of the zinc finger families. However, novel domains also appear such as the zinc binding LIM domains, some of which were shown to change their intracellular location upon redox regulation. The functional importance of these domains is further underlined by their involvement in various diseases that alter cysteine residues with a critical role in this type of regulation. Moreover, another large group of predicted RSCDPs corresponds to extracellular domains found only in multicellular organisms. The redox regulation of extracellular proteins could be important for their transfer to the extracellular matrix, and assembly in their target location. Thus, mutations of such critical, redox-regulated cysteine residues can lead to devastating diseases (Table 2). Coupling of potentially redox-sensitive thiols and structural plasticity is very well characterized in proteins transferred from cytosol to other organelles, such as mitochondria and peroxisome. Thus, the similar mechanism might be crucial for the biogenesis of extracellular proteins, including receptors. In addition to biogenesis, receptors could utilize redox-regulated regions for sensing changes in redox status and mediate related signaling events<u>[60] [62,63]</u>. Our results suggest that the human and drosophila proteomes share many common redox-sensitive structural switching domains, corresponding to similar functions. However, the abundance of these domains is further elevated in *Homo sapiens*, with the most drastic increase observed in the case of the C2H2 type zinc finger domains.

The predicted redox-sensitive structural switches were associated with several biological processes and functions, many of which have already been linked to changes in redox state. One important group includes transcription factors, many of which contain zinc finger domains[84]. In yeast, it has been shown that modulation of the redox status induces reversible changes in the translational machinery and controls protein synthesis[40]. Examples of redox-regulated proteins that rely on conditional disorder are involved in mitochondrial transport and assembly. In mammalians, redox signaling via alteration of redox status of specific protein thiols has been recognized as a major contributor to diverse key processes, such as stem cell proliferation[85], vertebrate embryonic development[86], neuronal development[87], and blood coagulation[88]. Redox regulation is also an emerging theme in cell-cell and cell-ECM interactions[89], tying together integrin signaling, angiogenesis, and various diseases. All of these processes are present only in higher order eukaryotes, and the heavy involvement of redox-sensitive conditionally disordered proteins is in line with the evolutionary trends identified (Figure 3).

Altogether, these results reinforce the importance of redox regulation in key biological processes. However, the analysis of human genetic variation also highlights that mutation of cysteine residues critical for such regulation are correlated with different pathologies (Table 2). This analysis shows that such disease-related mutations are found in proteins with different functions and different cellular localization. It is tempting to speculate, that some of these proteins might be involved in sensing or maintaining redox homeostasis in the related organelles. However, the understanding of the structural mechanisms underlying the development of these diseases have already served with potential therapeutic options. CADASIL commonly arises via NOTCH3 cysteine mutations, and artificially introduced exon-skipping removes the affected EGF-like domains, and can restore receptor functionality in vitro[90]. The large scale identification of similar redox-sensitive protein regions, together with pathway/mutation analysis can serve as new targets for therapeutic intervention. In addition to endogenous modulations leading to diseases, RSCDPs and redox regulation play critical roles in host-pathogen interactions as well. The role of redox-sensitive proteins in interactions between viruses and eukaryotic hosts is known for several examples, such as Hepatitis C virus [91], coxsackievirus B3 [92] and Epstein-Barr virus [93]. Viruses, in general, exploit the redox-state dependence of several signaling pathways of the host cell, and this has even been recognized as having therapeutic value [94]. Our results suggest that the use of redox-state dependent proteins can be a generic theme among viruses targeting multicellular hosts. Redox-state modulation is an emerging theme in the host-pathogen interactions of single-cell eukaryotic pathogens, such as trypanosoma, leishmania, malaria and others. Parasite survival and its successful proliferation depends on the capacity of the invading parasite to cope with the oxidation blast of the host immune system. Therefore an effective redox regulation mechanism is a crucial component for parasite survival which is found and characterized for many unicellular pathogens, including *T. congolense* and *P. falciparum*. Our analysis suggests that these pathogens contain significantly large fractions of proteins with redox-sensitive intrinsically disordered regions (Figure 3A). Further analysis of these proteins might have therapeutic advances. In order to choose potential candidates for experimental analysis, our method can be coupled with the recently developed experimental tools, which enable real-time mapping of redox states in the parasite[95].

In conclusion, the results presented in this study show that many protein regions rely on metal binding disulfide bonds for stability, which are potentially redox-regulated, and involve structural transitions from an ordered state to a disordered state, or vice versa. The abundance of these types of regions dramatically increases from yeast to human, and involves a wide range of processes related to multicellularity, signaling and regulation of transcription and translation.

## Acknowledgments

This work was supported by the “Lendület” grant from the Hungarian Academy of Sciences (LP2014-18) (Z.D., B.M., G.E.), OTKA grant (K108798) (Z.D., B.M., G.E.), the FIEK grant 5097/1/2018 (Z.D.), the EMBO|EuropaBio fellowship 7544 (B.M.), the Israel Science Foundation (1765/13) (D.R) and Human Frontier Science program (CDA00064/2014) (D.R). We thank Meytal Radzinski for the critical reading and editing of the manuscript.

## Data and Methods

For the prediction of redox-regulated conditionally disordered segments a slightly modified version of the IUPred2a method was used. The basic principle is the same as published [21]. We generate two prediction profiles with the default settings of the IUPred method, one for the native sequence corresponding to the state achieved through cysteine stabilization (redox-plus) and one calculated for a modified sequence with cysteines residues mutated to serine, corresponding to the state without cysteine stabilization (redox-minus). To initiate a redox-sensitive conditionally disordered region, the difference between the two profiles has to be larger than 0.3, while the minus profile is below 0.5 and the plus profile is above this threshold at a given position. This region is then extended until the difference between the two profiles becomes lower than 0.15, or the redox plus profile falls below 0.35. Two adjacent regions are combined if they are closer than 10 amino acids. Two minor modifications were introduced compared to the original web server implementation, aimed at decreasing potential overprediction of such regions. First, a predicted redox-sensitive conditionally disordered region must have at least 3 residues that meets the opening criterion. Second, the minimal length of the predicted region must be at least 15 residues. The current version of the program is available at: http://gerdos.web.elte.hu/data/iupred_redox_2.0.tar.gz

For the analyses of PDB structures, a non-redundant dataset of protein structures was downloaded from the PISCES web server filtered at 40% sequence similarity, resolution of at least 2 Å and maximum R factor of 0.25[23]. The dataset contained 1,4150 protein chains. 5 chains were omitted for technical reasons and for the remaining sequences the redox regions were predicted with the customized IUPred2A program. Disulfide bonds were identified from the structure using the distance criteria of 2.3 Å between the sulphur atoms of two cysteines. For the identification of cysteine clusters, all potential combinations of four non-disulfide bonded cysteine and histidine residues were considered within each structure. For the four selected residues, the approximate location of the metal center was calculated as the center of mass of the positions corresponding to the coordinates of SG atoms for cysteines and ND1 atom for histidines. A cysteine cluster was identified when the distance of each of the considered atoms from this center position was smaller than 4.2 Å. These criteria were selected to maximize the agreement with the provided PDB annotations.

Full proteomes were downloaded from the 23/08/2018 version of UniProt reference proteomes [24]. The customized IUPred2A was run on all protein sequences from all proteins, using the redox-sensitive option. A protein was considered to contain a redox-sensitive conditionally disordered region if IUPred2A returned at least one region. The f_redox_ value of a proteome was calculated as the number of proteins with predicted regions over the total number of regions in the proteome. In total, the analyzed data consisted of 192 viral, 436 archaeal, 608 bacterial, and 167 eukaryotic proteomes. The proteomes, together with the number of predicted redox-sensitive conditionally disordered proteins and f_redox_, are listed in the Supplementary material.

The set of human proteins containing predicted redox-sensitive conditionally disordered protein regions (RSCDP) was determined as described for all proteomes. The list of 5,644 proteins together with the 10,109 identified regions is listed in the Supplementary material.

Redox-sensitive conditionally disordered domains were identified by assessing overlapping regions predicted by IUPred2A, and domains predicted by Pfam [25] (release 31.0). A Pfam domain instance was considered to be redox-sensitive conditionally disordered if an IUPred2A region covers at least 5 of its residues and at least 20% of its total length. The number of thus identified redox-sensitive domains are listed in the Supplementary material for the *S. cerevisiae, D. melanogaster*, and human proteomes, containing 6,049, 13,775, and 21,050 proteins, respectively. In each case the reference proteome downloaded from UniProt were used.

Gene Ontology (GO) annotations for human proteins were taken from the 24/05/2018 version of the EBI GOA database [26]. Only those biological process terms were considered that are attached to at least one protein with at least one predicted redox-sensitive structural switch. In order to enable the analysis of fairly generic and high level processes, only those terms were kept which are reachable from the root term (biological_process) with at most four steps following only ‘is a’ or ‘part of’ relations. These terms were further filtered for ancestry, and if two terms were in ancestor/child relationship, only the child term was kept. The resulting set of GO terms were manually split into four groups based on the level of biological process they describe. This yielded the GO molecular slim, containing 61 terms describing basic molecular processes; the GO network slim, containing 77 terms describing pathway/network level processes; GO cell slim, with 103 terms describing cellular processes; and GO organism slim, with 183 terms describing organism level biological processes. The enrichment of each term among redox-sensitive conditionally disordered proteins was assessed comparing the occurrence of the term in the RSCDP set and the human proteome. 1,000 sets of 5,488 proteins were taken from the human proteome randomly, and were evaluated for term occurrences, yielding an average and standard error for expected occurrence. The enrichment of each term in RSCDP is expressed as the difference between the observed and expected occurrences in standard error deviations. The supplementary material shows the enrichment values for all terms in all four slims.

Missense variants annotated in human UniProtKB/Swiss-Prot entries were downloaded from the Humsavar website (https://www.uniprot.org/docs/humsavar). Only those mutations were considered that contained ‘Disease’ annotation as type of the variant and involved a cysteine either in the original or in the mutated form. Mutations located within redox-sensitive conditionally disordered regions were filtered out based on an overlap with predicted regions using the original sequence in cases when a cysteine residue was altered, or using the mutated sequence in case the mutation introduced a cysteine. The identified mutations are shown in the supplementary material. To concentrate on cases that are likely to be directly associated with alteration of a redox-regulated conditionally disordered region, proteins were sorted according to the ratio of mutations within predicted regions versus all mutations within the corresponding gene.

